# Do genetic differences in growth thermal reaction norms maintain genetic variation in timing of diapause induction?

**DOI:** 10.1101/849059

**Authors:** Erlend I. F. Fossen, Joost A. M. Raeymaekers, Sigurd Einum

## Abstract

1. An optimal timing for diapause induction through the sexual production of dormant propagules is expected in populations of annual organisms. Yet, experimental work typically finds high within-population genetic variation in the sexual production of such propagules. Thus, high genetic variation in timing for diapause induction should be a common feature of annual organisms.
2. Here, we hypothesize that genetic variation in the propensity to produce diapause propagules, *P*_*d*_, is maintained by a genotype-by-environment interaction in growth performance, where fast-growing genotypes within an environment should delay diapause relative to slow-growing genotypes. From this, we derive two predictions. First, if there is ecological crossover in growth performance, the genetic correlation of *P*_*d*_ between environments should be negative. Second, the correlation between absolute plasticities of growth and *P*_*d*_ should be negative.
3. We tested these predictions by quantifying ephippia production in genotypes of a population of the facultative sexual cladoceran *Daphnia magna* at two temperatures. The population biomass at the onset of ephippia production was used as a measure of *P*_*d*_, whereas somatic growth rate was used to quantify growth. Plasticity for both measurements was derived from thermal reaction norms.
4. Our results did not support either prediction, as neither the genetic correlation of *P*_*d*_ between environments, nor the correlation between absolute plasticity of growth and *P*_*d*_ were found to be significant.
5. Our results suggest that genetic variation in the timing of diapause is not maintained by genetic differences in thermal growth reaction norms. We propose as an alternative hypothesis that if there is across year variation in how stochastically the environment deteriorates, fluctuating selection may favor genotypes with different *P*_*d*_ between years.

## Introduction

The timing of reproduction is a crucial life history trait (Stearns, 2000). Many annual organisms, for which the end of the growing season results in death, produce dormant propagules that enable the survival of the population through harsh environmental conditions (e.g. winter or dry seasons). Examples of these dormant propagules are seeds in plants and diapause eggs in animals like killifish (Murphy and Collier, 1997), cladocerans (Frey, 1960), rotifers (García-Roger et al., 2017), copepods (Holm et al., 2018) and some insects (e.g. silkworms, Tirelli, 1946). For these organisms, the timing of the switch from somatic growth to the production of diapause propagules has strong fitness consequences, and there is an optimal timing of reproduction (Cohen, 1976). Delaying onset of reproduction allows for more time to grow and higher reproductive output per reproductive event, but comes at the cost of reducing the number of reproductive events or risking reproductive failure (Furness et al., 2015; Weis et al., 2014). Organisms with facultative parthenogenesis, which can switch between asexual and sexual reproduction (e.g. cladocerans [Gerber et al., 2018; Taylor and Gabriel, 1993], rotifers [García-Roger et al., 2017; Serra et al., 2008] and aphids [Simon et al., 2002]), represent a special case. For these, the optimal timing involves a trade-off between investing in biomass through repeated events of clonal reproduction vs. ensuring production of diapause eggs before the growing season ends.

Genetic variation in the timing of flowering and subsequent seed production is commonly observed in plants (e.g. Bourion et al., 2002; Franks et al., 2007; Hara and Ohsawa, 2013). Similarly, for facultative parthenogenetic animals which typically switch from asexual to sexual reproduction when environmental conditions deteriorate, there is genetic variation in the propensity to produce diapause propagules (*P*_*d*_) in a given environment, suggesting that genotypes vary in their environmental cue thresholds (e.g. Deng, 1996; Gilbert, 2002; Roulin et al., 2015; Yampolsky, 1992). In a seasonal environment, such variation in cue thresholds should translate into variation in timing of the switch. Thus, given the predicted optimal switch time within a population, explaining the maintenance of such genetic variation in *P*_*d*_ is a challenge. One hypothesis is that different genotypes are adapted to different environmental conditions that occur during different parts of the season, and thus show different responses to environmental gradients in their ability to accumulate biomass (i.e. genotype-by-environment [G×E] interactions in somatic growth or clonal reproduction, e.g. Carvalho, 1987). If this G×E interaction is strong, it can generate an “ecological crossover” where different genotypes are superior in different environments (Ellner and Hairston Jnr, 1994; Gillespie and Turelli, 1989; Higginson and Reader, 2009; Turelli and Barton, 2004). This should lead to corresponding differences in *P*_*d*_ across environments. For example, a genotype with low somatic growth rate in cold environments would benefit from having a higher *P*_*d*_ when exposed to a low temperature, compared to a genotype that is able to maintain a high somatic growth at that low temperature. Similar patterns have been observed for other genetically correlated traits, where these show corresponding G×E interactions (e.g. Mills et al., 2007; Prati and Schmid, 2000; Stinchcombe et al., 2004). For example, Prati and Schmid (2000) showed a genetic tradeoff between flowering and rooting (i.e. between sexual reproduction and clonal growth), where flowering showed a G×E interaction that corresponded with a G×E interaction in rooting. Thus, according to this hypothesis, two predictions can be derived regarding the patterns of genetic variance in *P*_*d*_ (Fig. 1). First, if there is ecological crossover in the ability to accumulate biomass (Fig. 1A), and *P*_*d*_ is quantified on either side of the environmental condition where crossing occurs, there should be a negative genetic correlation between *P*_*d*_ across environments (Fig. 1C). Second, there should be a negative genetic correlation between the absolute plasticity in *P*_*d*_ and the absolute plasticity in growth performance (Fig. 1D). The absolute plasticity is here defined as the slope of the trait reaction norm. The second prediction follows from the fact that, for a given genotype, the magnitude and direction of change in *P*_*d*_ across environments should depend on the corresponding change in growth performance.

**Figure 1:**
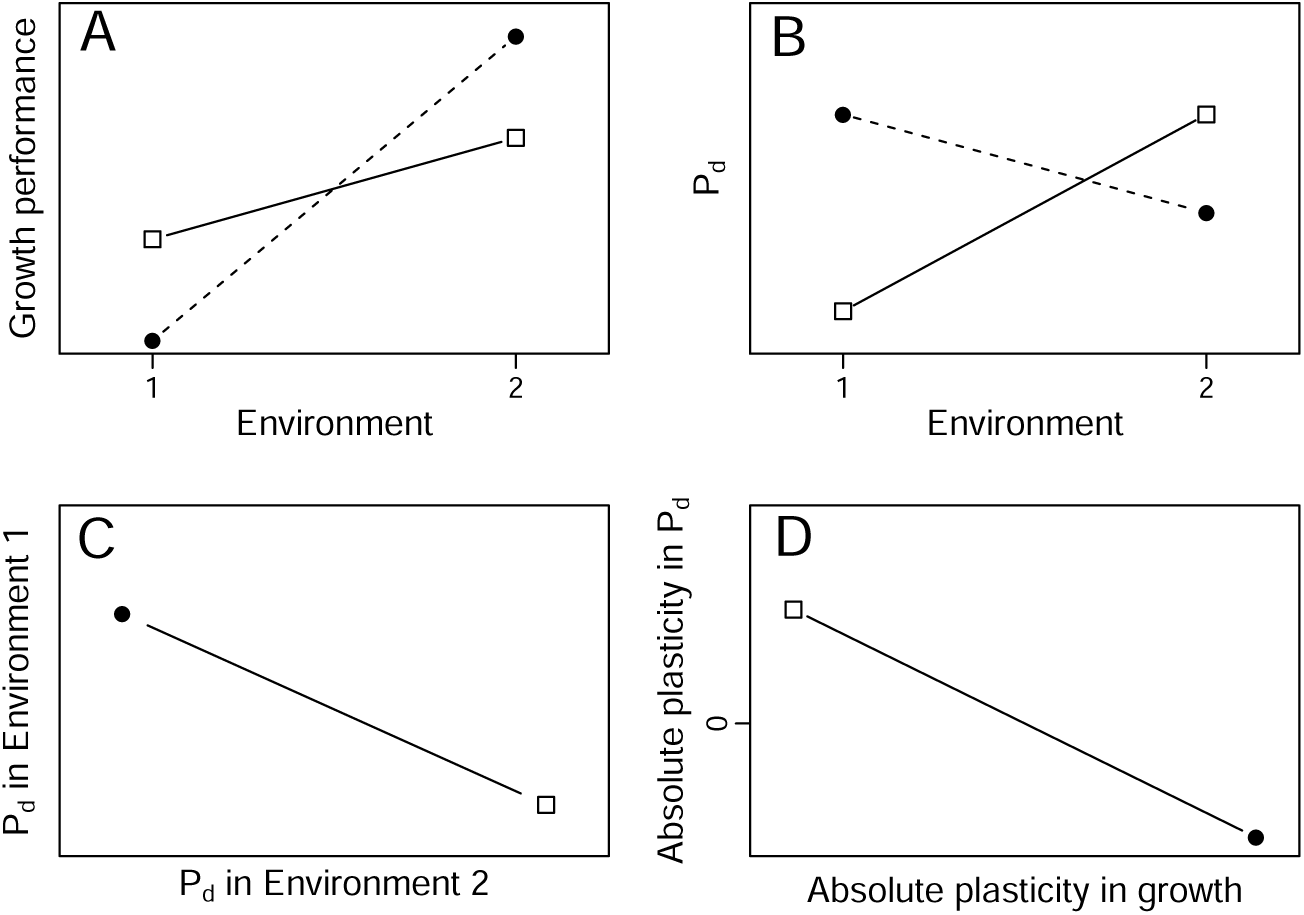
Predictions following the hypothesis that variation in reaction norms of growth performance can maintain genetic variation in the propensity to produce diapause propagules *P*_*d*_. If there is a strong G×E interaction with ecological crossover in growth performance (**A**, two genotypes represented by different symbols), reaction norms for *P*_*d*_ are expected to also show ecological crossover (**B**). Consequently, a negative genetic correlation is predicted between *P*_*d*_ in the two environments (**C**). Furthermore, we predict a negative genetic correlation between the absolute plasticity in these traits, where the absolute plasticity is defined as the slope of the trait reaction norm (**D**). This follows from the hypothesis that genotypes that experience a stronger decrease in growth performance when going from environment 2 to environment 1 should also have a higher increase in *P*_*d*_.

We tested these predictions by studying a population of the facultative sexual cladoceran *Daphnia magna*, a species that is known to harbor pronounced genetic variance in resting egg production (Roulin et al., 2015; Yampolsky, 1992). The population used originates from a pond at its northern distribution limit, where production of resting eggs during autumn is expected to be crucial for over-wintering fitness. Using a set of ten genetically distinct genotypes, we ran population growth experiments at different temperatures and quantified production of diapause propagules through resting egg counts. Population density is one important cue that *Daphnia* use to switch to resting egg production (Gyllström and Hansson, 2004; Kleiven et al., 1992). The population density required to trigger this switch was therefore used as a measure of *P*_*d*_ at the different temperatures. Furthermore, thermal reaction norms of somatic growth rates for the same clones (Fossen et al., 2018) were used to test for a genetic correlation between environmental cue thresholds and growth performance.

## Materials and methods

### Study animals and husbandry of stock cultures

Ephippia of *Daphnia magna* Straus, 1820, which contains up to two sexually produced resting eggs, were collected in November 2014 from the surface sediment of a shallow pond at Værøy Island (Sandtjønna, 1.0 ha, 67.687°N 12.672°E), northern Norway. This pond freezes over during winter and its mean daily water temperatures during summer fluctuates between 10-20°C (Fig. 2 in Fossen et al., 2018). Ten genotypes, hatched from separate ephippia in December 2014, were cultured separately under common garden conditions for 2-3 years (*>* 50 asexual generations). These genotypes, hereafter referred to as clones, vary genetically in thermal plasticity of life-history traits (Fossen et al., 2018). Daphnids were kept in 250 ml jars containing a modified ADaM medium (Klüttgen et al., 1994, SeO_2_ concentration reduced by 50%, seasalt increased to 1.23 g/L) at 17°C with a 16L:8D photoperiod, and the medium was changed weekly. Cultures contained five female adults per 250 ml jar and were fed three times a week with Shellfish Diet 1800 (Reed Mariculture Inc, Campbell, CA, USA) at a final algae concentration of 4×10^5^ cells ml^−1^.

**Figure 2:**
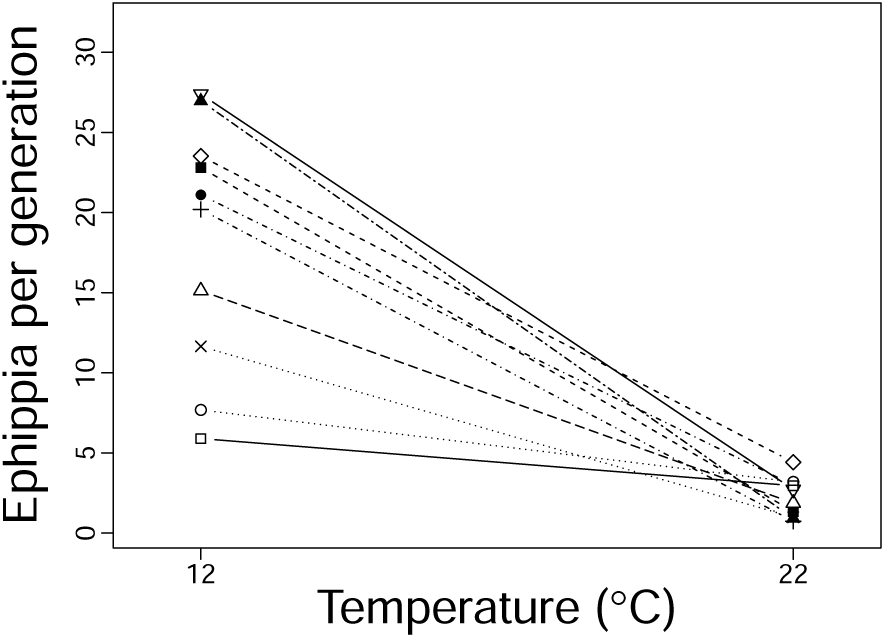
Ephippia production per generation across temperature for 10 clones of a population of *Daphnia magna*. Each point is the mean of a clone, represented by different symbols.

### Experimental design

*Daphnia* use several environmental cues to switch from growth to production of diapause propagules, including population density/food abundance, photoperiod, and temperature (e.g. Deng, 1996; Gyllström and Hansson, 2004; Kleiven et al., 1992; Slusarczyk and Rybicka, 2011). Of these, we kept photoperiod constant (16L:8D, representing autumn conditions in the population’s native environment), while experimentally manipulating temperature. To identify the temperatures under which the population initiate increased ephippia production, we conducted an exploratory experiment (May – December 2016, see Appendix S1 in supplementary information for details) using eight different temperature treatments (12, 15, 17, 19, 22, 24, 26 and 28°C), all with relatively low and similar mortality (Fossen et al., 2018). The results from this (Fig. S1) showed that the population produced ephippia at all temperatures, but that the total ephippia production was particularly high below 17°C, with little variation among the higher temperatures. Based on this we conducted a second experiment (January – June 2017, where experimental populations were started across 23 dates) using two temperature treatments (12 and 22°C) that were 5°C below and above the temperature that triggered the change in mean ephippia production, and which enabled testing our predictions.

The second experiment largely followed the same procedure as the first experiment (Appendix S1). Clones were kept in replicated lines at their experimental temperatures for two or more asexual generations prior to experiments to ensure acclimation. A single female juvenile (<24h old from the second clutch) was used to initiate each replicate experimental population. All temperature treatments (acclimation and experiment) were given by placing 250 ml jars in climate cabinets (IPP 260plus, Memmert, Germany). Throughout the acclimation and experimental period, animals were fed temperature-specific food concentrations every second day (concentrations 10^5^ cells ml^−1^: 12°C, 2.00; 22°C, 3.24) and the medium was refreshed every 8 days at 12 and every 4 days at 22°C. This food regime represents *ad libitum* concentrations when the population size is low (unpublished data), and ensures equal starting conditions across temperatures. The same food regime was used in Fossen et al. (2018) to obtain somatic growth rates, thus making the current study directly comparable to Fossen et al. (2018).

From the start of the second experiment until the second clutch had been born, each population was checked daily for offspring and the date and number of offspring was recorded. We obtained the exact timing of the onset of ephippia production by checking each population daily until the shedding of the first ephippium. At the onset of ephippia production, and after the second clutch was born, we recorded a six seconds video of each population at every second medium change (camera: Panasonic DMC-TZ25, Full HD, 1920×1080 pixels). For the video, all animals of a population were transferred to a transparent glass baking dish, which was put on top of a light table to obtain a high contrast between animals and the background. These videos were later used to estimate population sizes and body size distributions over time using the R-package *trackdem* v. 0.3.1 (Bruijning et al., 2018). Videos made by us (N = 153) under the same conditions as described above were used by Bruijning et al. (2018) to evaluate the accuracy of *trackdem*. This analysis revealed highly accurate and unbiased estimates of the population size (Fig. 3 in Bruijning et al., 2018). *Trackdem* uses automated particle tracking to keep track of the number of moving particles (here: individual daphnids) of potentially different size, in addition to outputting the size of the particles (number of pixels). By taking videos of animals of a known body mass (based on a known carapax length [CL, mm]) – dry mass [DM, mg] relationship: DM = 0.00535 · CL^2.72^, Yashchenko et al., 2016) we estimated the dry mass of each animal from its particle size (DM = −0.00635 + 0.00100×particle size, n = 21, R^2^ = 0.953). This allowed us to estimate total population size, the number of adults (animals *>* 0.075 mg dry mass) and the total biomass at any time a video was recorded. The number of shed ephippia was counted and stored during each medium change. The populations were stopped on day 128 at 12°C and on day 56 at 22°C, based on the results from the first experiment (Appendix S1). This corresponds to an intermediate time after peak population density where ephippia production also peaked in the first experiment. A total of 163 populations were studied in the second experiment, with 7-9 replicates per clone per temperature (Table S1).

**Figure 3:**
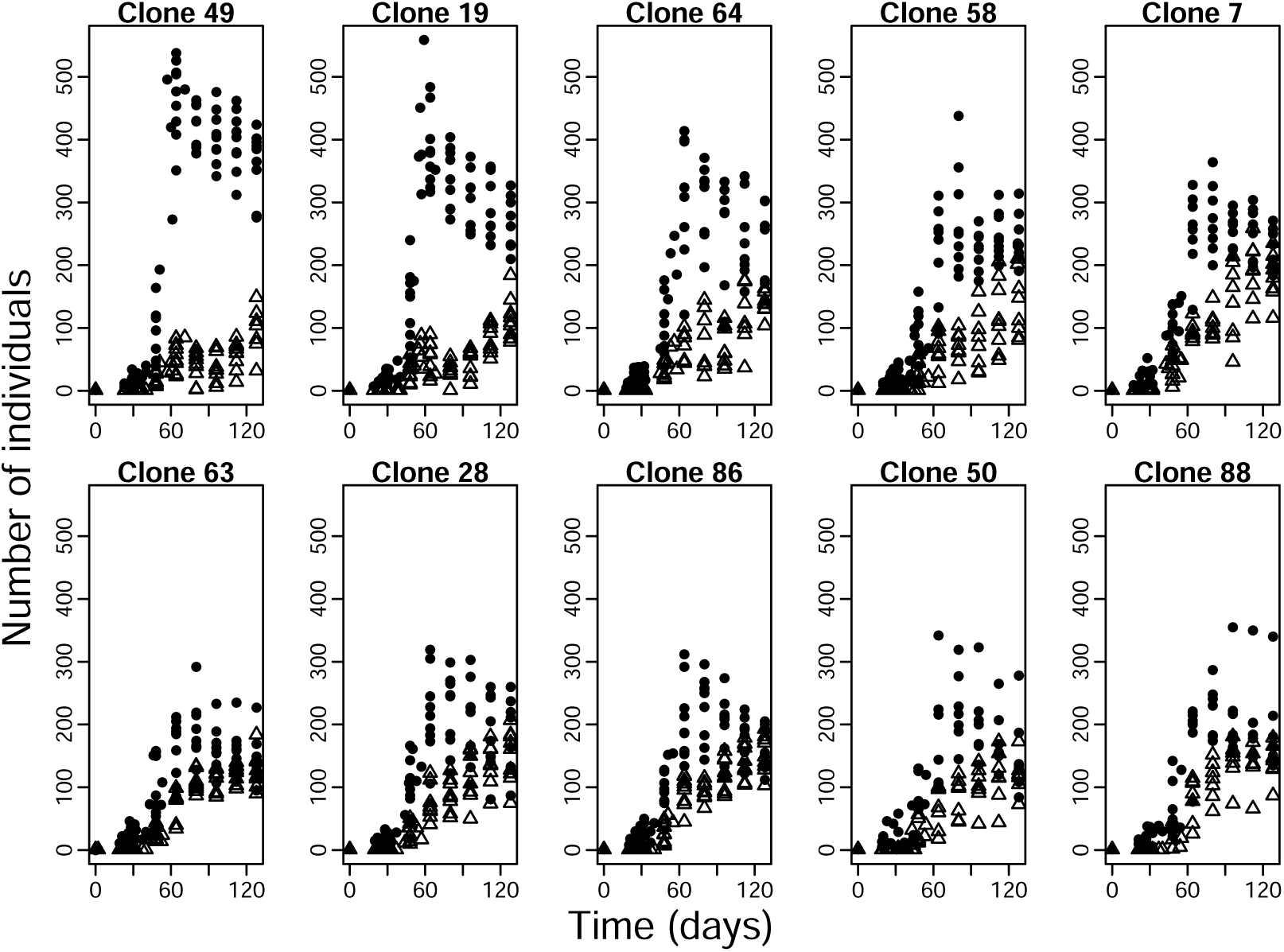
Association between ephippia production (panels sorted by increasing ephippia production from top left to bottom right) and age structure in 10 clones of *Daphnia magna* at 12°C. Filled circles represent the total population size, triangles the number of adults. There are multiple observations per time point because 7-9 replicates were used per clone.

### Trait measurements

#### Ephippia production

Ephippia production in *D. magna* involves production of resting eggs that require fertilization by males to be viable. Thus, for a given clone, producing few ephippia could potentially be compensated by predominately producing males (that fertilize receptive females from other clones), resulting in a negative correlation between production of males and production of ephippia. In the presence of such alternative reproductive tactics, a high production of males could be an alternative way to achieve genetic contribution to the diapause stage at the population level. Our design that kept clones separately could then potentially introduce noise when using ephippia production as a measure of *P*_*d*_. However, a previous study showed no such correlation among clones within populations of *D. magna*, and rather a positive correlation among populations (Roulin et al., 2015). Furthermore, we tested for this possibility in our own population by calculating the proportion of males of each population at the end of the first experiment. Although the proportion of males (mean [SD] = 0.025 [0.037]) differed significantly among clones (logistic GLM with clone as fixed effect, p < 0.001), there were no significant genetic correlations between the proportion of males and total ephippia production at any temperature (range: [-0.24, 0.23], p *>* 0.5 for all temperatures). This is a common finding for *Daphnia* (e.g. Lampert et al., 2012; Roulin et al., 2015; Yampolsky, 1992). Finally, the observation that an approximately 50:50 sex ratio is attained in populations under strong stimuli by production of alternating male and female clutches within females (Hobæk and Larsson, 1990), suggest that the presence of alternative reproductive tactics is unlikely. Thus, we only consider ephippia production in further analyses.

To get a measurement of ephippia production that is comparable across temperatures, we calculated ephippia produced per generation for each population as follows: 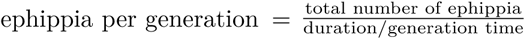, where duration is the duration of the time series, and the average time to first clutch of each temperature was used as a proxy for temperature-specific generation time.

#### Estimating P_d_

Population density and food availability are known cues used by *Daphnia* to switch from growth to ephippia production (Gyllström and Hansson, 2004; Kleiven et al., 1992). Thus, population biomass, which should better represent food availability than population density, should also be a valid cue for the switch. Thus, population biomass at the onset of ephippia production was used as a measure of *P*_*d*_ (i.e. a high population biomass required to trigger the switch to ephippia production represents a low *P*_*d*_). To make this comparable across temperatures (which differed in food rations) we divided this measure with the temperature specific food rations. While this may not completely control for differences in food ration, we note that all populations became food limited when reaching high population density and produced ephippia. Consequently, *P*_*d*_ could be estimated for each population and the pattern of genetic variance within temperatures should not be affected by not perfectly controlling for food ration across temperatures. Although we only estimated this for the second experiment where we had biomass measurements, a correlated measure (population density at the onset of ephippia production, Fig. S2A, B) was positively genetically correlated across experiments (Fig. S2C, D), indicating that this measurement is repeatable across experiments.

#### Growth performance

We used estimates of juvenile somatic growth rate from the same clones as used in this study as a measure of growth performance (Fossen et al., 2018). This trait has been shown to be a good proxy for the intrinsic rate of population increase in *D. magna* (Lampert and Trubetskova, 1996). An ecological crossover in somatic growth rate was found by Fossen et al. (2018), where reaction norms of somatic growth rate crossed at 14°C (Fig. S3). Clone-specific slope estimates of growth performance across temperatures were used to test our second prediction (Fig. 1).

### Statistical analyses

All analyses were conducted in R v.3.3.1 (R Core Team, 2014).

#### Total ephippia production – temperature response, genetic variance and links with age structure and P_d_

We used linear mixed-effect models with the package lme4 (v. 1.1-7, Bates et al., 2015) to test for genetic variance in total ephippia production across the two temperatures. Ephippia per generation was the response variable, and temperature was included as a categorical predictor variable. Clone and start date (of experimental populations) were used as random effects. Clone could affect the variance of the temperature treatments (i.e. clonal variance could vary with temperature) or the overall intercept, whereas start date was assumed to only affect the overall intercept. Akaike information criterion corrected for small sample sizes (AICc) were used in model selection. Full models with different random effect were first compared using restricted maximum likelihood (REML), and then the best random effect structure was used when comparing models with different fixed effects using maximum likelihood. Lastly, parameter estimates were obtained from the best model fitted with REML. Pseudo-R^2^ values were calculated as the squared correlation coefficient between fitted values from the model and observed values.

To explore how ephippia production was associated with population dynamics, we plotted how the age structure of populations changed over time. Using the mean proportion of adults over the entire time series for each clone, we estimated the genetic correlation between adult proportion and ephippia production. We also tested if having a smaller *P*_*d*_ resulted in lower total ephippia production. This was done by calculating genetic correlations between the two traits within each of the two temperatures, using temperature specific clonal means.

#### Prediction 1: G×E interactions and a genetic correlation between P_d_ across environments

To estimate genetic variance and test for G×E interactions in *P*_*d*_, we used mixed effect models with biomass per food abundance at the onset of ephippia production as the response variable. The same models and model procedures were applied as for ephippia production (see above). We then proceeded with testing for a negative genetic correlation between *P*_*d*_ across the two temperatures (as in Fig. 1C). We note that even if the estimated biomass per food abundance does not completely control for differences in food ration across temperatures, the genetic variation and order of clones within temperatures should not change and thus neither should the genetic correlation across temperatures.

#### Prediction 2: genetic correlation between the absolute plasticity of P_d_ and of growth performance

Our second prediction was that there should be a negative genetic correlation between the absolute plasticity in *P*_*d*_ and the absolute plasticity in growth performance (Fig. 1D). We quantified absolute plasticity in *P*_*d*_ as the negative of the slope (i.e. slope multiplied with −1) between biomass per food abundance at the onset of ephippia production and temperature, such that a clone with a large positive slope-value requires a relatively stronger population-density cue at low temperatures than at high temperatures. As for prediction 1, since genetic variance within temperatures should not be affected by us not completely controlling for differences in food ration across temperatures, neither should the genetic correlation between absolute plasticities be affected (even though the reaction norm slopes may change).

#### Evolutionary potential

To get an estimate of the population’s evolutionary potential that is comparable across traits, populations and species, the broad sense evolvability (clonal variance/mean^2^) was calculated within temperatures (Hansen et al., 2011, 2003; Houle, 1992). This was done for both ephippia production and *P*_*d*_. Evolvability is a measurement of the expected percentage change in a trait per generation under a unit strength of selection, and is (in contrast to heritability) independent from environmental variance (Hansen et al., 2011).

## Results

### Total ephippia production – temperature response, genetic variance and links with age structure and P_d_

Ephippia production was about eight times larger at 12 than 22°C (Fig. 2). We found a significant G×E interaction in ephippia production (Table S2), with higher evolvability at 12°C (V_clone_ = 51.34, evolvability = 14.27%) than at 22°C (V_clone_ = 0.08, evolvability = 2.74%). Variance due to start date was 12.67, the residual variance was 19.49, and the model pseudo-R^2^ was 0.86. Furthermore, the variation in ephippia production among clones was highly repeatable across experiments at 12°C (Fig. S1B), but not at 22°C (Fig. S1C).

Ephippia production at 12°C increased for all populations after about 60 days (Fig. S4), shortly after peak population density (Fig. 3). Further-more, ephippia production at 12°C was genetically correlated with age structure in the population (r = 0.93, t_8_ = 7.16, p < 0.001), where clones that produced many ephippia also tended to be dominated by adults compared to low producing clones that were more dominated by juveniles (Fig. 3). At 12°C but not at 22°C, *P*_*d*_ was positively genetically correlated with ephippia production (Fig. S5).

### Prediction 1: G×E interactions and a genetic correlation between across P_d_environments

*P*_*d*_ showed a significant G×E interaction with temperature, with some clones having a particularly low propensity to produce diapause propagules at low temperatures (Fig. 4A, Table S2). The overall evolvability was higher at 12°C (V_clone_ = 1664.4, evolvability = 35.45%) than at 22°C (V_clone_ = 2.2, evolvability = 0.08%). The model pseudo-R^2^ was 0.75, variance due to start date was 117.0 and the residual variance was 414.5. However, in contrast to the prediction of a negative genetic correlation between *P*_*d*_ at 12 vs 22°C, the genetic correlation of the trait across temperature was not significant (Fig. 4B, r = 0.41, t_8_ = 1.26, p = 0.243).

**Figure 4:**
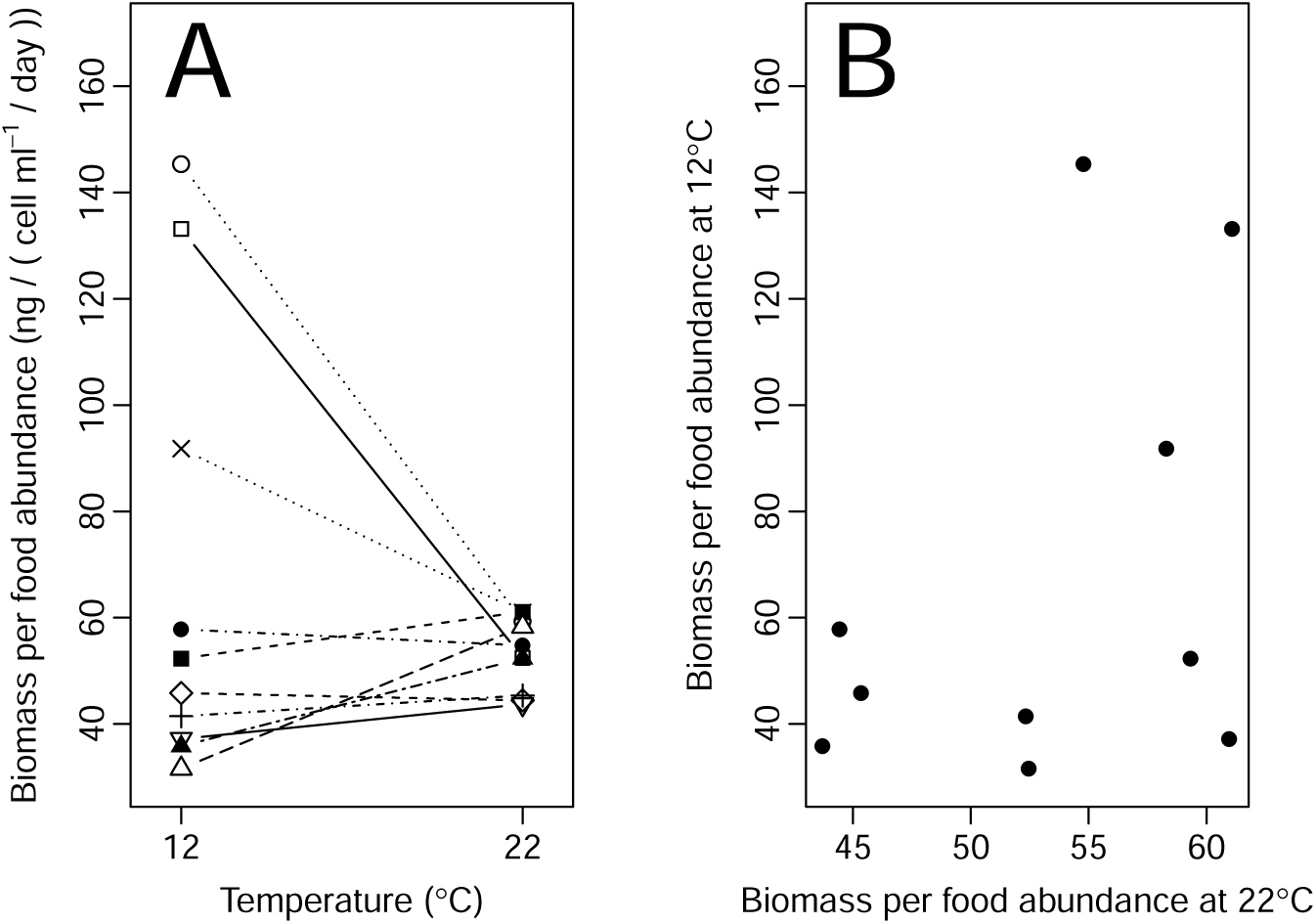
Propensity to produce diapause propagules (*P*_*d*_) for 10 clones of a population of *Daphnia magna. P*_*d*_ was measured as population biomass per food abundance at the onset of ephippia production, where a low biomass represents a high *P*_*d*_. There is a significant G×E interaction (**A**), but no significant genetic correlation across temperatures (**B**, r = 0.41, t_8_ = 1.26, p = 0.243).

### Prediction 2: genetic correlation between the absolute plasticity of P_d_ and of growth performance

While we expected absolute plasticity in *P*_*d*_ to decrease with increasing absolute plasticity in somatic growth rate, the relationship between the traits was not significant (Fig. 5, r = 0.29, t_8_ = −0.855, p = 0.417).

**Figure 5:**
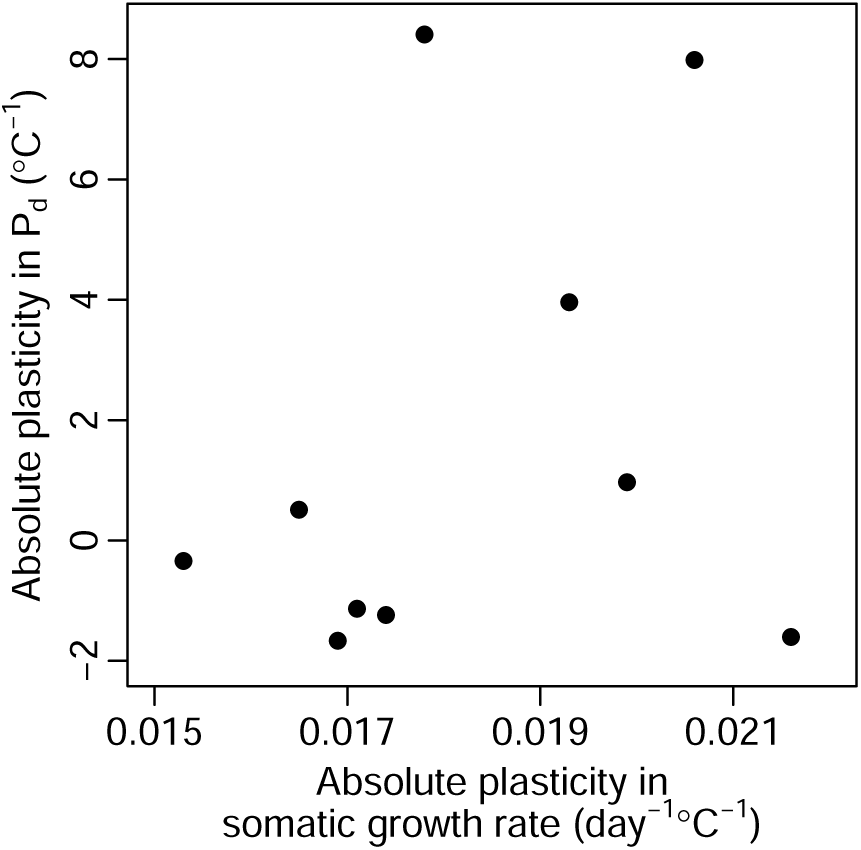
No significant genetic correlation (r = 0.29, t_8_= −0.855, p = 0.417) between thermal plasticity in the propensity to produce diapause propagules (*P*_*d*_) and a growth performance trait (somatic growth rate). Each point shows the absolute plasticity of the traits, representing the mean change in the trait per °C for a given clone of *Daphnia magna*. The absolute plasticity of growth rate was measured as the slope of the traits reaction norm, whereas absolute plasticity in *P*_*d*_ was measured as the negative of the slope between biomass per food abundance at the onset of ephippia production and temperature.

## Discussion

To test if genetic differences in thermal growth reaction norms can maintain genetic variation in the timing of diapause induction, we ran population growth experiments using clones from a single population of *Daphnia magna*. We quantified relationships between ephippia (resting egg) production, the propensity to produce diapause propagules (*P*_*d*_), and a growth performance trait (somatic growth rate) across two temperatures. Our results showed that there was high genetic variance in *P*_*d*_, and *P*_*d*_ was positively genetically correlated with total ephippia production. Both traits showed higher evolvability (*>* 10% at 12°C) than what is usually found in wild populations (about 1% for life history traits, Hansen et al., 2011), particularly at 12°C. The observed differences among clones at 12°C were also highly consistent with results from a separate exploratory experiment (Fig. S1, S2). Having found such large differences among clones in *P*_*d*_, we tested a hypothesis derived from the observation that genotypes within the same population can be adapted to different seasons (e.g. Carvalho, 1987; Grosberg, 1988), i.e. that genotypes should match genotype-specific environmental variation in *P*_*d*_ with their corresponding variation in their asexual production. We tested two predictions derived from this hypothesis (Fig. 1). First, we predicted a negative genetic correlation between *P*_*d*_ across a temperature range that encompass ecological crossover in asexual production. Second, we predicted a negative genetic correlation between absolute thermal plasticity in somatic growth rate and absolute thermal plasticity in *P*_*d*_. Our results did not support either of the predictions, indicating that genetic differences in growth thermal reaction norms do not maintain genetic variation in *P*_*d*_.

Although our predictions should enable testing if genetic differences in growth thermal reaction norms maintain genetic variation in *P*_*d*_, certain conditions may have prevented us from detecting the predicted outcomes. For instance, if the temperature treatments were too similar, this could prevent detection of a negative correlation between *P*_*d*_ across temperatures. However, the fact that thermal reaction norms for somatic growth rate cross at a temperature intermediate to our two experimental temperatures, and that ephippia production showed large differences at the tested temperatures, suggests that the temperature treatments were sufficiently different to detect any genetic correlation between *P*_*d*_ across temperatures. Alternatively, if juvenile somatic growth rate was a poor measure of asexual population growth, a genetic correlation between absolute plasticity in somatic growth and absolute plasticity in *P*_*d*_ would not be predicted. This does however seem unlikely considering that this measure of somatic growth rate has previously been strongly linked to the intrinsic rate of population increase in our study species (Lampert and Trubetskova, 1996). Overall, for both predictions to fail simultaneously, two independent conditions would need to occur (too similar test-environments and using an inappropriate trait as a measure of growth performance), which seems unlikely.

An alternative explanation for the maintenance of high genetic variation in *P*_*d*_ is that there is strong fluctuating selection (across years) on this trait. This can occur if there is variation across years in to what extent temperature regimes deviate from a smooth seasonal trend of spring increase and subsequent fall decrease. In years with high levels of stochastic deviation from such a trend, short periods of low temperatures are likely to be followed by subsequent increases that allows for continued high asexual growth under low population densities. This will lead to lost opportunities for genotypes that have a high *P*_*d*_, and hence rapidly switch to resting egg production in the absence of strong competition (i.e. at low population density). Selection should then favor “optimist” clones that have a low *P*_*d*_ and require stronger population density cues to commence resting egg production during such years. At the other extreme, for years with no stochasticity a decline in temperature is always followed by a further decline. In this situation, selection should favor more “pessimistic” clones that do not wait and see whether temperature conditions improve following a decline, but that have a high *P*_*d*_ and rapidly engage in resting egg production when exposed to a certain decline in temperature. In our experimental data, such “pessimist-optimist” strategies were associated with age structure (Fig. 3), where more “optimistic” clones (with a low *P*_*d*_ and lower total ephippia production) became more dominated by juveniles than more “pessimistic” clones. This association with age structure also clearly illustrates the tradeoff between producing asexual offspring and producing sexual resting eggs. Such maintenance of genetic variation due to temporally fluctuating selection is similar to other cases where the optimum trait value fluctuates over time (e.g. timing of diapause in copepods, Hairston and Dillon, 1990; morph color and pattern in stick insects, Nosil et al., 2018). These “pessimist-optimist” strategies may also explain the low genetic variance at 22°C compared to at 12°C. If temperature conditions are favorable, all clones should invest in growth until their growth rate is constrained to a certain level due to high competition, resulting in all clones requiring a similar and high population density threshold to initiate resting egg production.

Declines in temperature have been suggested as a cue for the onset of winter and for inducing diapause in freshwater zooplankton in temperate areas (Gyllström and Hansson, 2004; Slusarczyk and Rybicka, 2011). Our finding of ephippia production being much higher at 12 than at 22°C supports this. Yet, the median biomass per food abundance needed to induce ephippia production was similar at the two temperatures. This may seem surprising, but can be explained by the fact that ectotherms (including our *Daphnia* clones, Fossen et al., 2019) at higher temperatures have higher metabolic rates and therefore also higher food demands. This explanation is supported by the observation that populations were able to maintain a much higher mean biomass per food abundance after the first ephippium had been produced at 12°C (mean ± SE = 162.0 ± 3.3 ng/[cell ml^−1^ day^−1^]) than that at 22°C (mean ± SE = 65.6 ± 1.5).

In this study, we tested if a strong G×E interaction in growth performance can maintain genetic variation in the propensity to produce diapause propagules. Despite finding high genetic variance in *P*_*d*_, and using a growth performance trait (somatic growth rate) that showed a strong G×E interaction, we found no support for this hypothesis. *P*_*d*_ showed no genetic correlation across temperature (prediction 1), and there was no genetic correlation between absolute plasticity in growth performance and absolute plasticity in *P*_*d*_ (prediction 2). We propose that genetic variance in *P*_*d*_ can be maintained by fluctuating selection favoring either “pessimist” (i.e. expressing high values of *P*_*d*_ when experiencing declines in temperature) or “optimist” genotypes, depending on how stochastically the environment deteriorates within a certain year. It can be expected that the stochasticity of environmental deterioration will vary from year to year, particularly for temperate populations. Consequently, “pessimist-optimist” strategies may explain genetic variation in timing of diapause induction for a wide range of annual organisms.

## Acknowledgments

Financial support was provided by the Research Council of Norway, FRIPRO programme, project ‘Eco-evolutionary dynamics of thermal reaction norms’ (Project 230482), and partly by the Research Council of Norway through its Centres of Excellence funding scheme, project number 223257/F50 and the Norwegian University of Science and Technology (NTNU). We thank V. Yashchenko, H.-K. Lakka, M. Jeannot, V. Parry, A. Vold, and others for help with data collection and culture maintenance.

## Author Contributions

EIFF, JAMR and SE contributed with conceptual ideas and developed study design. EIFF and JAMR collected the data. EIFF conducted statistical analyses with input from JAMR and SE. EIFF wrote the initial draft of the manuscript. JAMR and SE contributed in revisions.

## Data archival location

Data will be archived in Dryad Digital Repository.

## SUPPORTING INFORMATION

## Appendix S1: Experiment 1

### Experimental design

To identify the temperatures under which the population initiate increased ephippia production, we conducted an exploratory experiment (May – December 2016) using eight different temperature treatments (12, 15, 17, 19, 22, 24, 26 and 28°C). Clones were kept in replicated lines at their experimental temperatures for two or more asexual generations prior to experiments to ensure acclimation. A single female juvenile (<24h old from the second clutch) was used to initiate each replicate experimental population, and each of these replicates (within clone and temperature) originated from unique grandmothers to reduce maternal effects when estimating genetic effects. All temperature treatments (acclimation and experiment) were given by placing 250 ml jars in climate cabinets (IPP 260plus, Memmert, Germany). Throughout the acclimation and experimental period, animals were fed temperature-specific food concentrations every second day (concentrations 105 cells ml^−1^: 12°C, 2.00; 15°C, 2.38; 17°C, 2.62; 19°C, 2.88; 22°C, 3.24; 24°C, 3.50; 26°C, 3.76; 28°C, 4.00) and the medium was refreshed every 8 days at 12 and 15°C, every 6 days at 17 and 19°C, every 4 days at 22 and 24°C, and every 2 days at 26 and 28°C. The food supply represents ad libitum concentrations when the population size is low (unpublished data). Each experimental temperature treatment started approximately at the same time to ensure overlap in time within the experiment. From the onset of the experiment until the second clutch had been born, each population was checked daily for offspring and the date and number of offspring was recorded. After the second clutch was born and until the population size started to decrease, we estimated the population size by manually counting the number of individuals during every medium change. After the population started to decrease, we only estimated the population size every second medium change. The number of shed ephippia was counted and stored during each medium change. The populations were stopped after 225 days at 12°C, 201 days at 15°C, 162 days at 17°C, 126 days at 19°C, 120 days at 22°C, 108 days at 24 and 26°C, and 102 days at 28°C. These days correspond to the time point at which populations started to stabilize in population size, based on when a temperature-specific estimate of carrying capacity (based on Ricker’s model, Ricker, 1954) changed by less than 3% from one population count to the next. All remaining *Daphnia* were collected and each population was stored separately on ethanol after the populations were stopped. The number of replicates per clone per temperature was unbalanced with a median of four replicates at 17 and 19°C, three replicates at 12°C, two replicates at 22-28°C, and one replicate at 15°C (Table S1).

### Statistical analyses

To identify at which of eight temperatures increased ephippia production was initiated, we used ANOVA with post hoc Tukey tests. Total ephippia production was the response variable and temperature was used as a categorical predictor variable.

### Results

Total ephippia production was 4 to 5 times higher at 12 and 15°C than at ≥ 17°C (Fig. S1A, adjusted p < 0.001 for all two-way comparisons), and the 12 and 15°C treatments were not significantly different from each other (adj. p = 0.996). None of the ≥ 17°C treatments were significantly different from each other (p *>* 0.995 for all comparisons).

---

**Table S1:** Sample sizes per clone by temperature for each experiment.

| Experiment | Temperature<br>(°C) | Clone |  |  |  |  |  |  |  |  |  |
| --- | --- | --- | --- | --- | --- | --- | --- | --- | --- | --- | --- |
|  |  | EF7 | EF19 | EF28 | EF49 | EF50 | EF58 | EF63 | EF64 | EF86 | EF88 |
| 1 | 12 | 0 | 5 | 2 | 3 | 0 | 3 | 4 | 3 | 4 | 6 |
|  | 15 | 1 | 1 | 1 | 1 | 1 | 1 | 1 | 0 | 1 | 1 |
|  | 17 | 4 | 3 | 3 | 4 | 4 | 3 | 4 | 3 | 4 | 4 |
|  | 19 | 4 | 3 | 3 | 4 | 4 | 3 | 4 | 4 | 3 | 4 |
|  | 22 | 2 | 2 | 2 | 2 | 2 | 2 | 2 | 2 | 2 | 2 |
|  | 24 | 2 | 4 | 2 | 2 | 2 | 2 | 2 | 3 | 2 | 2 |
|  | 26 | 2 | 2 | 2 | 2 | 2 | 2 | 2 | 2 | 2 | 2 |
|  | 28 | 2 | 2 | 2 | 2 | 2 | 2 | 2 | 2 | 2 | 2 |
| 2 | 12 | 8 | 9 | 9 | 9 | 7 | 9 | 9 | 8 | 9 | 7 |
|  | 22 | 8 | 8 | 8 | 8 | 8 | 8 | 8 | 8 | 7 | 8 |

**Table S2:**
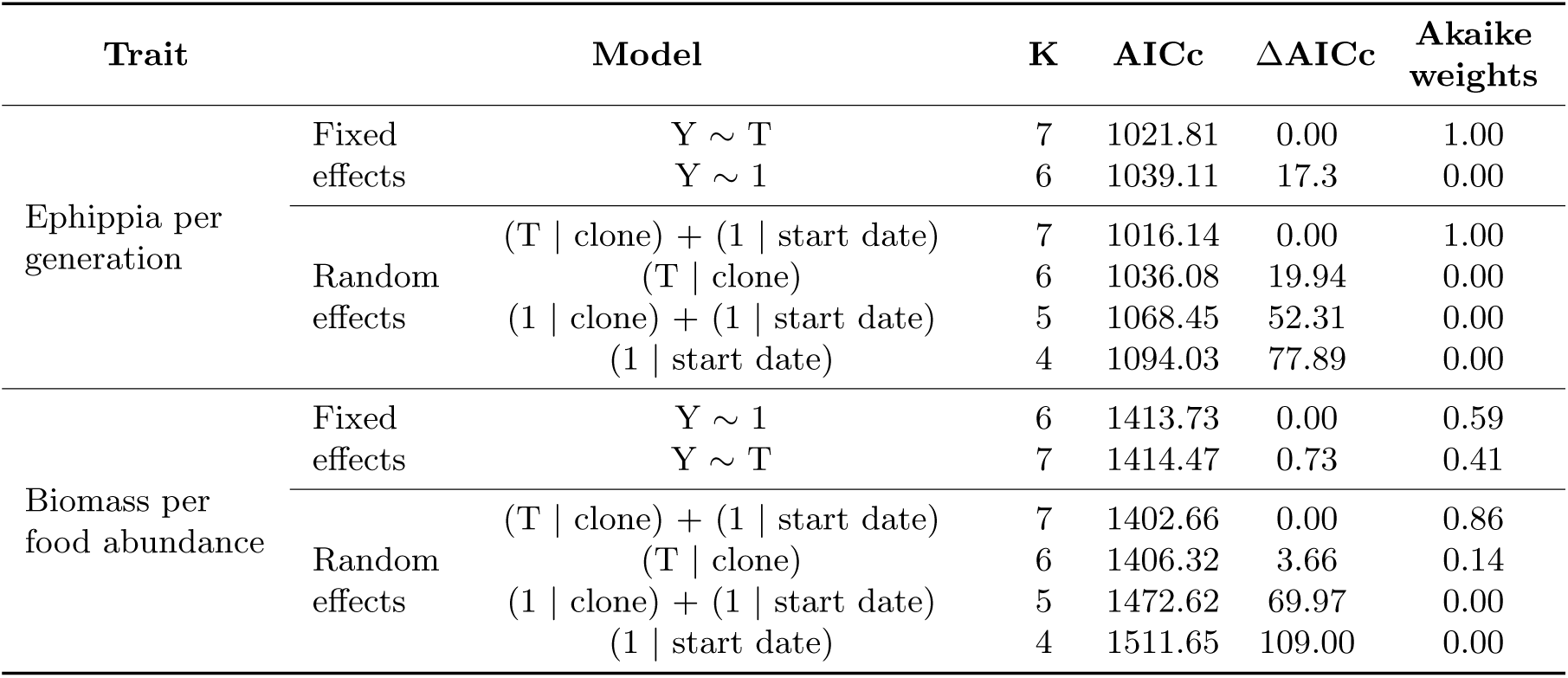
Model selection using AICc of candidate models for ephippia per generation and *P*_*d*_ (measured as biomass per food abundance (ng/[cell ml^−1^ day^−1^]) at the onset of ephippia production) for a population of *Daphnia magna*. Models sorted by ΔAICc. The best random effect structure was first determined with REML on models that included all listed fixed effects. Fixed effects were then compared with ML using the best random effect structure. T = temperature, used as a categorical variable; K = number of parameters.

**Figure S1.**
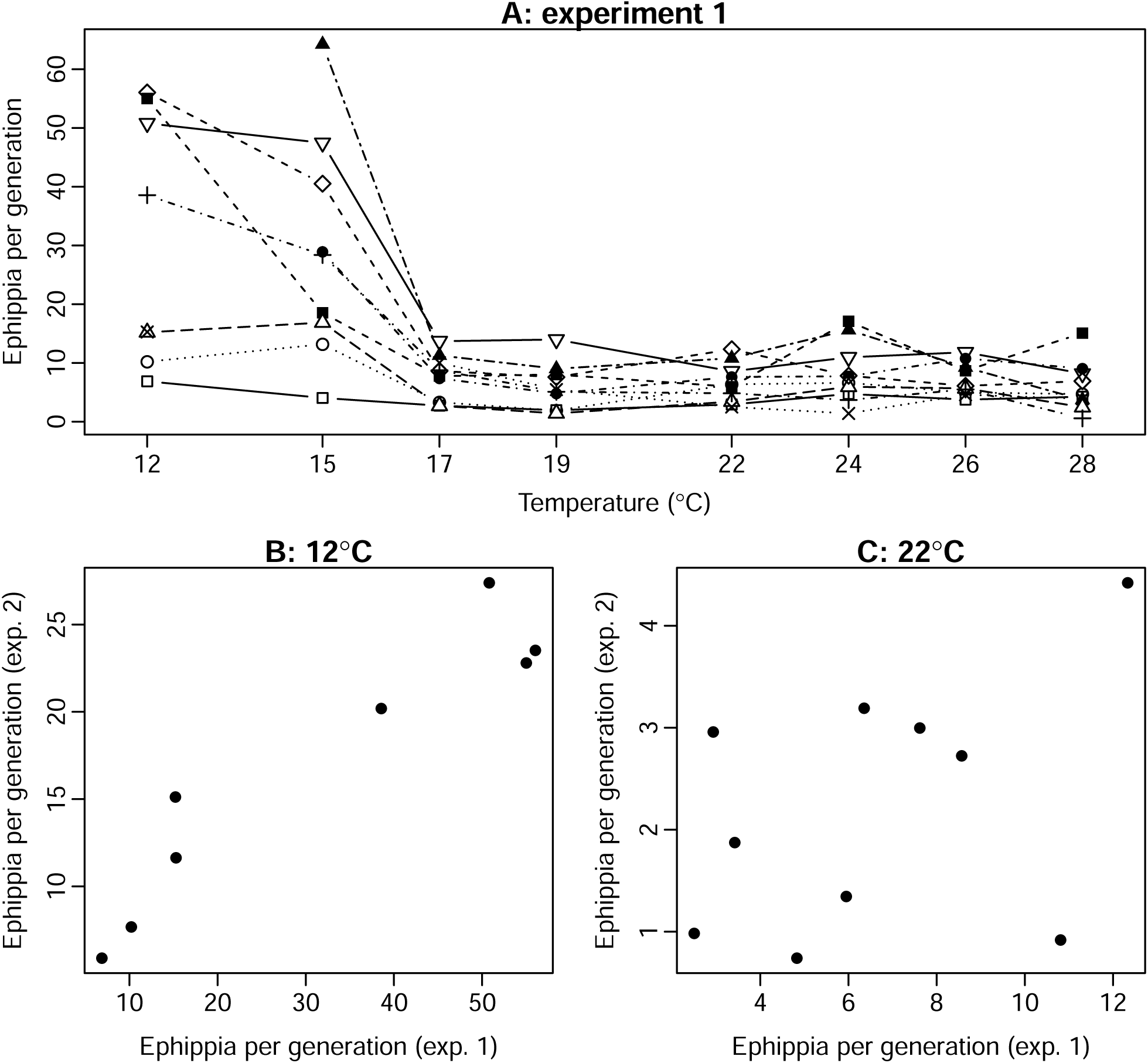
Ephippia production per generation across temperature in experiment 1. Each point is the mean of one of 10 clones of a population of *Daphnia magna*, represented by different symbols (**A**). **B**) High repeatability of ephippia production across experiments at 12°C (genetic correlation r = 0.94, t_6_ = 6.94, p < 0.001). C) Weak repeatability across experiments at 22°C (r = 0.42, t_8_ = 1.30, p = 0.229).

**Figure S2.**
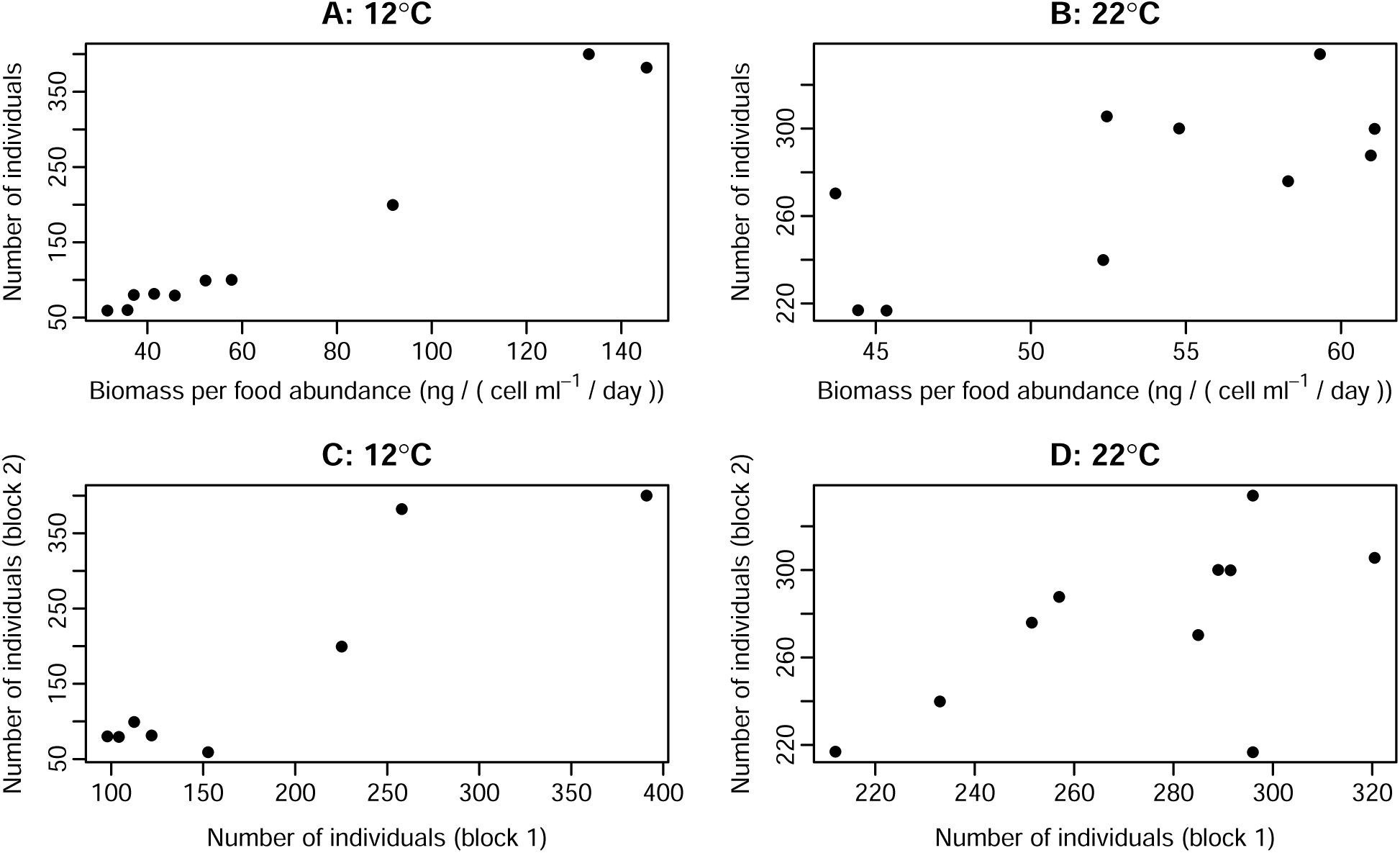
Genetic correlations between biomass at the onset of ephippia production and the number of individuals at the onset of ephippia production at 12°C (**A**, r = 0.99, t_8_ = 16.7, p < 0.001) and at 22°C (**B**, r = 0.71, t_8_ = 2.89, p = 0.020). Note that a high biomass represents a low *P*_*d*_. Each point is the mean of a clone of *Daphnia magna*. Number of individuals was highly genetically correlated across experiments at 12°C (**C**, r = 0.92, t_6_ = 5.76, p = 0.001), and weakly correlated at 22°C (**D**, r = 0.59, t_8_ = 2.08, p = 0.072).

**Figure S3.**
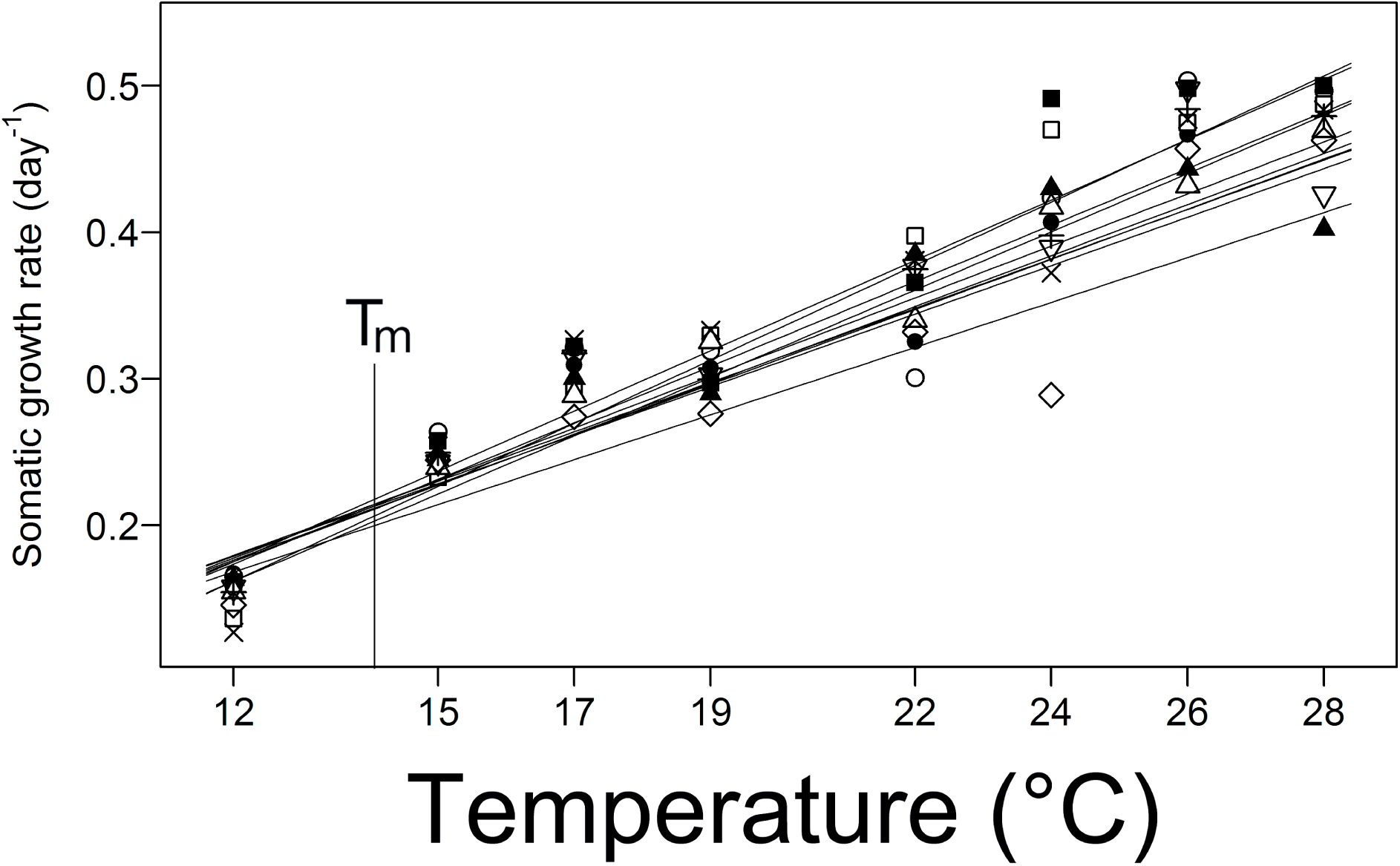
Thermal reaction norms of somatic growth rate for the same clones of *Daphnia magna* as used in this study (Fig. 1b in Fossen et al., 2018). Somatic growth rate shows ecological crossover with a zone of canalization (T_*m*_) at 14°C, where reaction norms cross. See Fossen et al. (2018) for more details.

**Figure S4.**
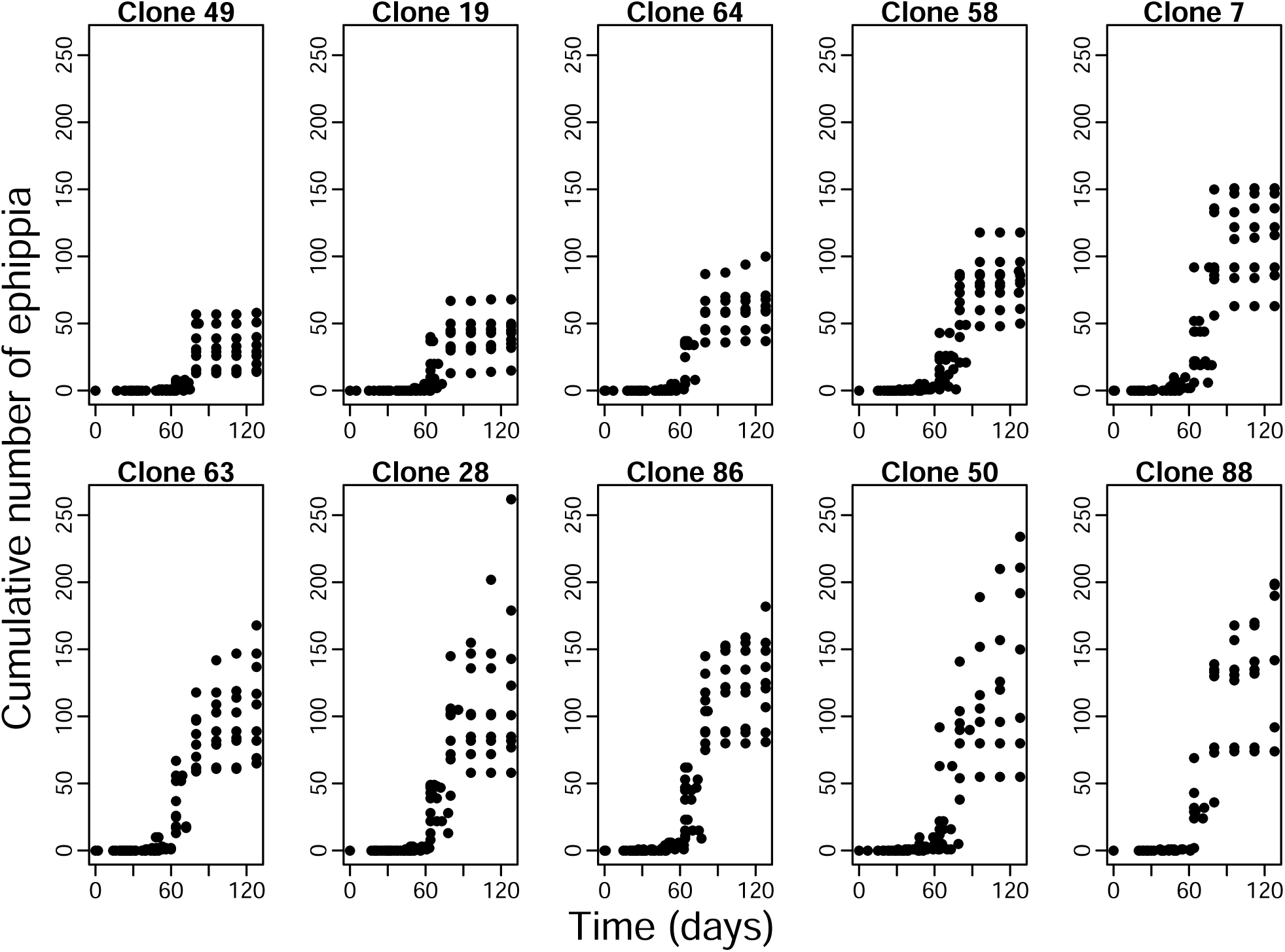
Cumulative number of ephippia over time at 12°C for 10 clones of *Daphnia magna*. Multiple points per time point because 7-9 replicates were used per clone. Panels are sorted by increasing mean overall ephippia production from top left to bottom right (same order as Fig. 3).

**Figure S5.**
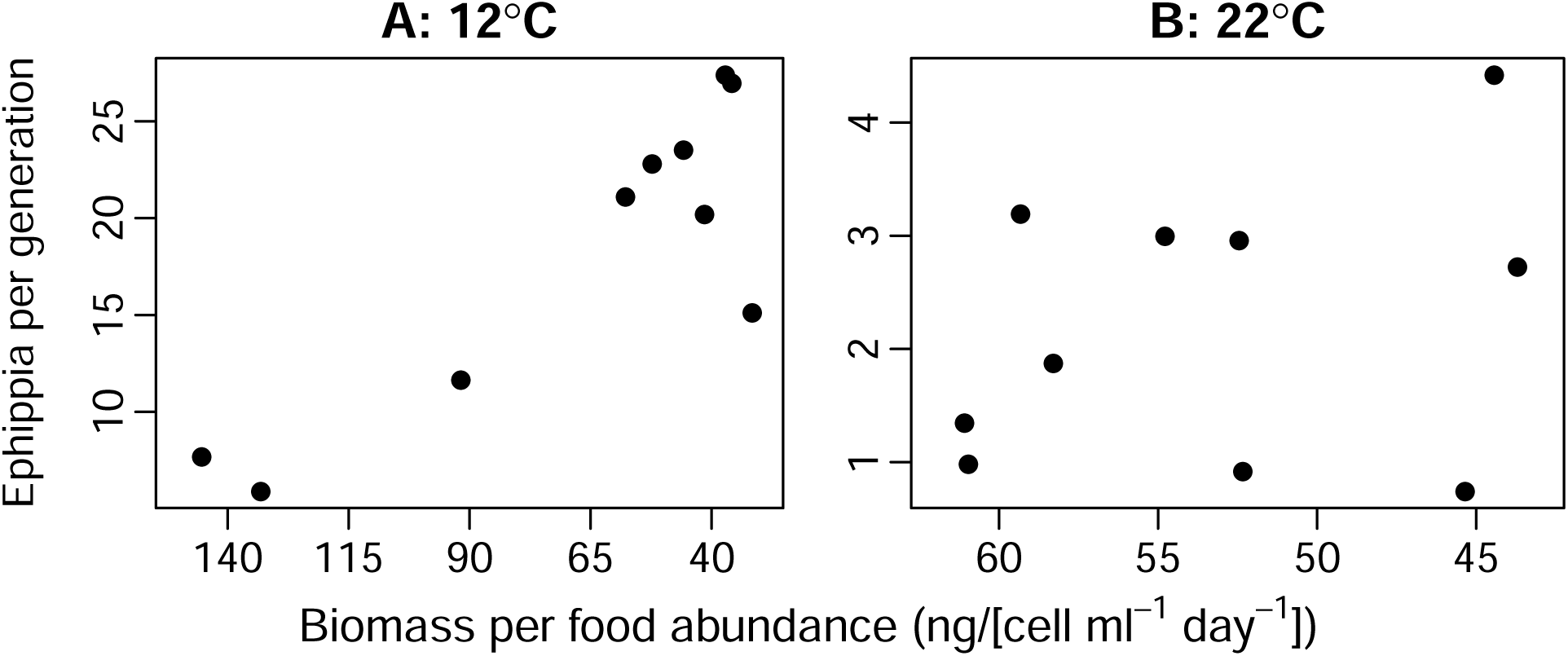
Genetic correlations between propensity to produce diapause propagules (*P*_*d*_) and ephippia production 10 clones of *Daphnia magna. P*_*d*_ was measured as the population biomass per food abundance at the onset of ephippia production, where a low biomass represents a high *P*_*d*_. **A**) Strong negative correlation between biomass and ephippia at 12°C (r = −0.86, t_8_ = −4.79, p = 0.001), corresponding to a positive genetic correlation between *P*_*d*_ and ephippia production. **B**) No significant correlation between biomass and ephippia at 22°C (r = −0.31, t_8_ = −0.908, p = 0.390).

